# Comparing DNA extraction methods for successful PacBio HiFi sequencing: a case study of the freshwater mussel *Anodonta anatina* (Bivalvia: Unionidae)

**DOI:** 10.1101/2025.11.26.690746

**Authors:** Marco Giulio, Urs Lergster, Philine G. D. Feulner, Alexandra A.-T. Weber

**Affiliations:** Department of Aquatic Ecology, Swiss Federal Institute of Aquatic Science and Technology (Eawag), Dübendorf, Switzerland; Department of Fish Ecology & Evolution, Swiss Federal Institute of Aquatic Science and Technology (Eawag), Kastanienbaum, Switzerland; Institute of Ecology and Evolution, University of Bern, Bern, Switzerland

**Keywords:** Mollusc genomics, inhibitors, reference genome

## Abstract

High-quality reference genomes are increasingly recognized as essential resources in biodiversity genomics and conservation. However, successful DNA extraction and long-read sequencing remain highly organism-dependent. Molluscs, a diverse phylum of invertebrates, pose particular challenges due to the presence of inhibitory compounds and the difficulty of obtaining high-molecular-weight DNA, often necessitating careful optimization of extraction protocols. Here, we present a case study on the freshwater mussel *Anodonta anatina* (Bivalvia: Unionidae), evaluating two preservation methods, six DNA extraction protocols, and two post-extraction clean-up steps for their effects on DNA quality and PacBio HiFi sequencing yield from foot tissue of a single individual. The PacBio Nanobind and CTAB protocols produced high-quality DNA from fresh tissue but performed poorly on flash-frozen tissue. Post-extraction clean-up generally degraded DNA and did not improve sequencing yield. Unexpectedly, the column-based Omega Mollusc Kit, although not designed for high-molecular-weight DNA, performed better than the PacBio-recommended Nanobind kit and the manual CTAB method on flash-frozen tissue. It generated high DNA quantity and purity, sufficient integrity for HiFi sequencing, and appeared to remove contaminants effectively. While the resulting DNA may be too fragmented for ultra-long read sequencing, the Omega Mollusc Kit offers a practical, cost-effective first approach for testing DNA extraction and PacBio sequencing in flash-frozen *A. anatina* foot tissue. When fresh tissue is available, Nanobind or CTAB were the best-performing options in our comparison. Overall, our results provide a practical starting point for protocol selection, while acknowledging that validation across other mollusc species, tissue types, and preservation methods remains important. This strategy could reduce the need for extensive protocol optimization and facilitate future mollusc genomics efforts.

## Introduction

The generation of reference genomes plays a crucial role in the conservation and management of biodiversity, providing a foundation for understanding species’ evolutionary histories, adaptive potentials, and responses to environmental pressures (Formenti et al., 2022; Paez et al., 2022; Theissinger et al., 2023). Large-scale initiatives such as the Earth BioGenome Project (Lewin et al., 2018, 2022) with dedicated local centers, for instance the European Reference Genome Atlas (Mc Cartney et al., 2024), have emerged to coordinate the global effort to sequence and catalog genomes from all eukaryotic species. These projects are especially important for species-rich groups such as molluscs, one of the most diverse animal phyla with around 87,000 described extant species, with gastropods and bivalves being the two most species-rich classes (MolluscaBase eds. 2025).

Despite the high number of mollusc species, genomic resources for this phylum lag behind other species-rich taxa (Gomes-dos-Santos et al., 2020; Johansen and Davison, 2025). The reasons for this gap are multifaceted. While some challenges are attributed to genome complexity (e.g. heterozygosity and repeat content) (Halstead-Nussloch et al., 2024; Chen et al., 2025), a more pervasive issue involves the co-extraction of compounds that inhibit DNA polymerase activity during sequencing. Molluscs are known to produce polyphenolic proteins, mucopolysaccharides, and other biochemical contaminants that complicate the extraction of high-quality DNA, hampering efforts to obtain accurate and complete genomic data (Adema, 2021).

Accordingly, numerous DNA extraction protocols for molluscs have been developed not only to increase DNA yield, but also to address different classes of co-extracted inhibitors. In particular, CTAB-based methods have often been adopted because CTAB, especially under high-salt conditions, helps separate DNA from polysaccharides and associated contaminants, whereas other modified precipitation or purification steps have been introduced for tissues especially rich in mucus and mucopolysaccharides (Sokolov, 2000; Pereira et al., 2011; Arseneau et al., 2017; Chakraborty et al., 2020). Nevertheless, ensuring sufficient DNA quantity, integrity, and purity for successful sequencing remains challenging, because contaminants are not always completely removed during extraction and can compromise downstream enzymatic reactions even when DNA yield is high (Arseneau et al., 2017; Adema, 2021). Commercial kits specifically designed for molluscs, such as the E.Z.N.A. Mollusc & Insect DNA Kit (Omega Bio-tek), have been developed to address these challenges. These kits typically produce DNA with good yield and purity, making them a popular choice for routine DNA extractions. However, their efficacy is supposedly limited when long-read sequencing technologies are required (e.g. to assemble reference genomes) (Jaudou et al., 2022). The frequent use of centrifugation and column-based purification steps in these kits typically results in DNA fragmentation, which is not optimal for techniques such as Oxford Nanopore or PacBio where long DNA fragments (typically >50 kb) are usually required for *de novo* genome sequencing and assembly (Arseneau et al., 2017; Angthong et al., 2020). In addition, these approaches generally produce relatively low yields of high-molecular-weight (HMW) DNA (Kalendar et al., 2023). Nevertheless, this kit was recently applied in the *Anodonta cygnea* genome project, where it enabled successful PacBio HiFi sequencing (Faust et al., 2025), suggesting that its utility for mollusc genome projects may be greater than previously assumed.

Despite adherence to established DNA quality control measures, such as achieving appropriate Nanodrop ratios, good DNA integrity, and quantity, as well as successful library preparation for long-read sequencing platforms like PacBio, the expected sequencing outputs are not always met. For example, cases have been reported where sequencing results yielded shorter reads and lower output than anticipated, despite following correct protocols (Mollusc Genomics Conference 2024, pers. comm.). In such cases, there may still be unknown contaminants co-purified with the DNA, which could potentially inhibit the performance of the high-fidelity polymerase, ultimately compromising the sequencing output. Unfortunately, such issues remain largely undetectable prior to sequencing, as both the DNA and libraries may appear high-quality based on standard QC metrics. As a result, incorporating additional post-extraction DNA clean-up steps may help eliminate residual contaminants and improve sequencing performance. Finally, it has been shown that tissue preservation method (e.g. flash freezing in liquid nitrogen and preservation at -80°C; preservation in ethanol 100%; or using fresh tissue directly) can also influence the outcome of DNA extractions (Alberola-Mora et al., 2025), with flash freezing in liquid nitrogen being the current gold standard for *de novo* reference genome projects (Reichel et al., 2026).

To date, systematic studies that compare long-read sequencing output from different preservation methods and DNA extraction protocols remain uncommon, with some recent exceptions (Angthong et al., 2020; Alberola-Mora et al., 2025). This is a significant gap in the context of large-scale genome initiatives where optimizing genomic resources for non-model organisms such as molluscs is critical. In this study, we address this gap by conducting a comparison of various factors that can influence DNA extraction and sequencing success in the freshwater mussel *Anodonta anatina* (Bivalvia: Unionidae), a species notorioulsy difficult to extract high-molecular weigth DNA from (Faust et al., 2025). Specifically, using foot tissue from the same individual, we: 1) Compare two different preservation methods (flash freezing in liquid nitrogen; fresh tissue); 2) Compare six DNA extraction protocols; 3) Evaluate the impact of two additional DNA clean-up steps applied before library preparation; and 4) Assess the sequencing output from five libraries sequenced on one full PacBio Revio cell. Our goal was to provide a comprehensive evaluation of the factors influencing genome sequencing, thereby offering practical guidance for improving genomic resource generation in freshwater mussels and, hopefully, more generally to other molluscs and other non-model organisms.

## Methods

### Sampling and molecular species identification

A collection permit (Nr. 24091) was obtained from Canton Zürich (Switzerland) and five adult individuals of the duck mussel *Anodonta anatina* were collected on the 5^th^ of August 2024 by snorkeling in Türlersee, Switzerland (GPS coordinates: 47.265602, 8.510178). They were brought back to the laboratory and kept alive in the Eawag experimental facility Aquatikum until processing. They were maintained in an open-water circuit at 19°C and fed twice a week using a mixture of lab-grown algae. Given the morphological plasticity of *Anodonta* mussels (Riccardi et al., 2020), species identification was initially based on shell morphology and subsequently confirmed by sequencing a COI barcode. Briefly, a small tissue biopsy was taken from the foot of each of the five individual mussels and DNA was extracted using the E.Z.N.A Tissue DNA kit (Omega Bio-tek) according to the manufacturer’s instructions. DNA integrity, quantity, and purity were assessed using NanoDrop Eight. A fragment of the COI mitochondrial gene was amplified with forward primer LCO22me2: 5’-GGTCAACAAAYCATAARGATATTGG-3’ and reverse primer HCO700dy2: 5’-TCAGGGTGACCAAAAAAYCA-3’ (Walker et al., 2006). PCR cycling conditions were: an initial step of 95°C for 15 min, followed by 40 cycles of 94°C for 30 s, 48°C for 40 s, 72°C for 60 s and then a final extension of 72°C for 10 min. Amplified PCR products were purified and Sanger sequenced by Microsynth AG, Balgach, Switzerland. The COI sequences of the five *Anodonta* individuals were compared to the NCBI GenBank database using the Basic Local Alignment Search Tool (BLAST) to confirm morphological species identification.

### Comparison of DNA extraction methods and quality control

All DNA extractions were performed using ∼30 mg of foot tissue from a single *Anodonta anatina* individual to eliminate inter-individual variability and isolate the effects of methodological factors. This design allowed us to focus on methodological rather than biological sources of variation, but did not allow inter-individual variation in extraction performance to be quantified. We tested the impact of two variables on DNA extraction success (Fig. 1): 1) Preservation method: fresh tissue vs. flash frozen in liquid nitrogen and subsequently preserved at -80°C; 2) DNA extraction method: six protocols, five of which being specifically designed to extract high-molecular weight (HMW) DNA (Table 1). For both preservation methods, tissues were processed immediately upon sampling: fresh foot tissue was transferred directly into lysis buffer after biopsy, and tissues intended for the frozen treatment were dissected and flash frozen in liquid nitrogen immediately after sacrifice. Each extraction condition was performed in triplicate, except for the GenomicTip and MegaLong protocols with frozen tissue, which were conducted in duplicate. All DNA extractions were conducted following the manufacturer’s instructions or using a dedicated protocol (Table 1). For the Nanobind and Omega kits, tissue homogeneisation was performed with a TissueRuptor II (QIAGEN). For the Omega kit, a single final elution step was conducted. DNA extractions using the six protocols were first performed on fresh foot tissue, as this allowed non-lethal sampling (small biopsy) and kept the individual alive. Then, the specimen was killed and remaining tissues (foot, gills, and whole body) were flash frozen in liquid nitrogen and stored at –80°C. Subsequent extractions from flash frozen foot tissue were conducted using the six DNA extraction methods after approximately three months at –80°C. Finally, to evaluate the effect of clean-up on DNA quality, we applied two commercial clean-up kits (Zymo and PowerClean) only to fresh-tissue extracts from the two top-performing column-free methods (CTAB and Nanobind), while limiting additional sample consumption. For each extraction method, the three replicate extracts were first pooled, and each pooled sample was then divided into three aliquots: one aliquot was cleaned with the Zymo kit, one with the PowerClean kit, and one was left untreated as a no-clean-up control.

**Table 1.** List of the DNA extraction methods and DNA clean-up kits used in this study. HMW: high-molecular weight.

| Kit or protocol name | Purpose | Short name | Kit price (CHF) | Number of preps | Price per prep (CHF) | Link |
| --- | --- | --- | --- | --- | --- | --- |
| PacBio Nanobind panDNA kit | HMW DNA extraction for animals | Nanobind | 315 | 24 | 13.1 | https://www.protocols.io/view/sanger-tree-of-life-hmw-dna-extraction-manual-moll-14egn36nyl5d/v1 |
| CTAB for molluscs | HMW DNA extraction for molluscs | CTAB | 237 | ~3000 | 0.08 | https://www.protocols.io/view/ctab-dna-extraction-protocol-for-mollusks-e6nvwjpr2lmk/v1 |
| Omega Bio-tek E.Z.N.A. Mollusc & Insect DNA Kit | DNA extraction for mollusc | Omega | 480 | 200 | 2.4 | https://omegabiotek.com/product/e-z-n-a-mollusc-dna-kit/ |
| Qiagen MagAttract | HMW DNA extraction | MagAttract | 350 | 48 | 7.3 | https://www.qiagen.com/us/products/discovery-and-translational-research/dna-rna-purification/dna-purification/genomic-dna/magattract-hmw-dna-kit-48 |
| Qiagen Genomic-tip 100G | HMW DNA extraction | GenomicTip | 422 | 25 | 16.9 | https://www.qiagen.com/us/products/discovery-and-translational-research/dna-rna-purification/dna-purification/genomic-dna/qiagen-genomic-tips?catno=10223 |
| G-Biosciences MegaLong | HMW DNA extraction | MegaLong | 768 | 25 | 30.7 | https://www.gbiosciences.com/MegaLong/MegaLong-359 |
| Qiagen DNeasy PowerClean Pro Cleanup Kit | DNA clean-up | Powerclean | 362 | 50 | 7.2 | https://www.qiagen.com/us/products/discovery-translational-research/dna-rna-purification/dna-purification/dna-clean-up/dneasy-powerclean-pro-cleanup-kit |
| ZYMO Genomic DNA Clean & Concentrator-10 | DNA clean-up | Zymo | 133 | 25 | 5.3 | https://zymoresearch.eu/collections/genomic-dna-clean-concentrator-gdcc/products/genomic-dna-clean-concentrator-25 |

**Figure 1:**
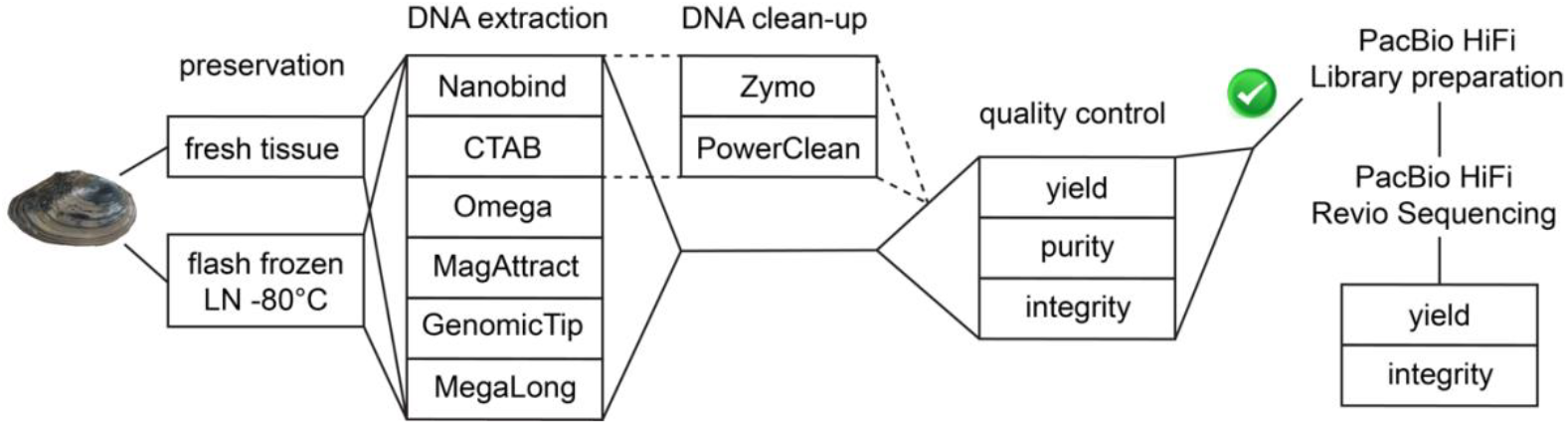
Experimental design for comparing DNA extraction methods and sequencing output in the freshwater mussel *Anodonta anatina*. All DNA extractions were performed from foot tissue of a single individual. DNA quality control at the DNA extraction laboratory was based on Qubit yield, NanoDrop purity, and TapeStation integrity; DNA integrity of selected samples was re-assessed at the sequencing center using a Fragment Analyzer prior to library preparation. LN: liquid nitrogen. See Table 1 for details regarding DNA extraction methods.

We recorded several metrics for DNA quality control: 1) Yield (Qubit dsDNA High Sensitivity kit); 2) Purity (Nanodrop 260/280 ratio); 3) Integrity (Agilent TapeStation gDNA ScreenTape). A DNA extraction was considered successful if it met all of the following thresholds: 1) Yield: >1 µg total DNA; 2) Purity: 260/280 ratio between 1.8 and 2.0; 3) Integrity: largest TapeStation peak >15 kb. 260/230 ratio was not used as a formal threshold because it was not consistently predictive of downstream sequencing success in our workflow.

### PacBio library preparation and sequencing

A total of 15 DNA extractions from eight treatments were brought to the Lausanne Genomic Technologies Facility (GTF) sequencing platform in Lausanne (Switzerland) for quality control, PacBio library preparation and sequencing (Table 2). DNA integrity was first assessed using a 5200 Fragment Analyzer (Agilent Technologies). Based on the quality thresholds described above, six DNA extractions representing six treatments were selected for PacBio library preparation. To avoid sequencing multiple extracts from the same treatment, only the best-performing extraction from each selected treatment was retained (CTAB_fresh1; CTAB_fresh_Zymo1; Nanobind_fresh1; Nanobind_fresh_Zymo1; Omega_frozen1; CTAB_frozen1; Table 2). DNA fragments below <10 kb were partially removed with a short-read eliminator (SRE) kit (PacBio). DNA of Omega_frozen1 sample was not sheared because it was already fragmented enough. For the five remaining DNA extractions, DNA was sheared in a Megaruptor 3 DNA shearing system (Diagenode) with a target mean fragment length of 15-20 kb. Fragments profile was assessed by capillary electrophoresis on a Fragment Analyzer using an HS large fragment 50 kb kit (Agilent Technologies). Repair and A-tailing of DNA fragments, SMRTbell adapters ligation, nuclease treatment and removal of fragments <3 kb were performed according to the SMRTbell prep kit 3.0 protocol (PacBio). Primer annealing and polymerase binding were performed using the PacBio Revio SPRQ polymerase kit including a DNA internal control according to the manufacturer’s protocol. Sequencing run was set up using SMRT Link v25.01. The six libraries were sequenced in HiFi mode in a PacBio Revio platform using Revio SPRQ Sequencing plate and one 25M ZMW SMRT cell with a 30-hour movie time and 2 hours of pre-extension time. Unfortunately, the initial run failed due to a technical issue with the PacBio instrument and had to be repeated. Due to the exhaustion of the CTAB_fresh_Zymo1 library, only five libraries were ultimately sequenced on a single 25M ZMW SMRT cell.

**Table 2.** Quality control summary of the 15 DNA extractions brought to the sequencing facility GTF. Extraction protocol, tissue preservation and potential clean-up steps are included in the sample name. All DNA extractions were performed using foot tissue from the same *A. anatina* individual. Technical replicates are numbered 1-3. The size detection limit of the genomic DNA Tapestation is 60,000 bp. Tapestation QC was performed in the extraction laboratory in Dübendorf. Fragment Analyzer QC was performed at the sequencing facility GTF in Lausanne. **(*)** Samples used for library preparation. ^1^ yield calculated from Qubit concentration.

| Sample name | Treatment | Yield<sup>1</sup><br>[ug] | Qubit<br>[ng/μl] | Nanodrop<br>[ng/μl] | A260/A280 | A260/A230 | Largest peak<br>Tapestation [bp] | Largest peak<br>Fragment<br>Analyzer [bp] |
| --- | --- | --- | --- | --- | --- | --- | --- | --- |
| (*) CTAB_fresh1 | 1 | 4.8 | 48.4 | 122.5 | 1.94 | 1.62 | >60,000 | 48,313 |
| (*) CTAB_fresh_Zymo1 | 2 | 1.9 | 41.6 | 53.7 | 1.88 | 3.07 | 51,900 | 47,003 |
| CTAB_fresh_Powerclean1 | 3 | 1.5 | 33 | 57.9 | 1.84 | 1.92 | 12,973 | 147 |
| (*) Nanobind_fresh1 | 4 | 7.5 | 74.6 | 159.8 | 1.86 | 2.24 | >60,000 | 47,751 |
| (*) Nanobind_fresh_Zymo1 | 5 | 1.6 | 36.2 | 40.4 | 1.76 | 3.16 | >60,000 | 40,640 |
| Nanobind_fresh_Powerclean1 | 6 | 0.6 | 13.4 | 16.3 | 1.92 | 1.04 | 17,352 | 8,557 |
| (*) Omega_frozen1 | 6 | 18.8 | 125 | 209.7 | 1.84 | 2.17 | 13,080 | 24,733 |
| Omega_frozen2 | 6 | 93.8 | 625 | 902.8 | 1.87 | 2.34 | 13,519 | 22,861 |
| Omega_frozen3 | 6 | 75.8 | 505 | 1,111.6 | 1.86 | 2.37 | 14,075 | 22,300 |
| Nanobind_frozen1 | 7 | 90 | 600 | 920.8 | 1.93 | 2.34 | 27,603 | 3,810 |
| Nanobind_frozen2 | 7 | 38.1 | 254 | 493.5 | 1.96 | 2.33 | >60,000 | 3,953 |
| Nanobind_frozen3 | 7 | 124.5 | 830 | 974.9 | 1.91 | 2.39 | 22,985 | >60,000 |
| (*) CTAB_frozen1 | 8 | 42.6 | 284 | 461.3 | 1.92 | 2.71 | 18,259 | 21,177 |
| CTAB_frozen2 | 8 | 41.6 | 277 | 457.5 | 1.93 | 2.72 | 10,900 | 84 |
| CTAB_frozen3 | 8 | 39.9 | 266 | 426.7 | 1.92 | 2.76 | 14,120 | 12,813 |

## Results

### Molecular species identification, comparison of DNA extraction performance and DNA clean-up

Using COI barcodes, all five analyzed individuals were assigned to *Anodonta anatina*, confirming the morphological species identification. A single representative individual (AE01) was subsequently used for all DNA extraction tests. Full QC metrics for all extractions are provided in Table S1. Three high-molecular weight (HMW) DNA extraction protocols— MagAttract, GenomicTip, and MegaLong—failed to produce DNA of sufficient quality, regardless of whether fresh or flash-frozen tissue was used. These protocols did not meet the minimum quality criteria for yield, purity, or integrity (Fig. 2; Table S1). In particular, the GenomicTip and MegaLong kits yielded almost no detectable DNA, and therefore none of these extracts were selected for sequencing.

**Figure 2.**
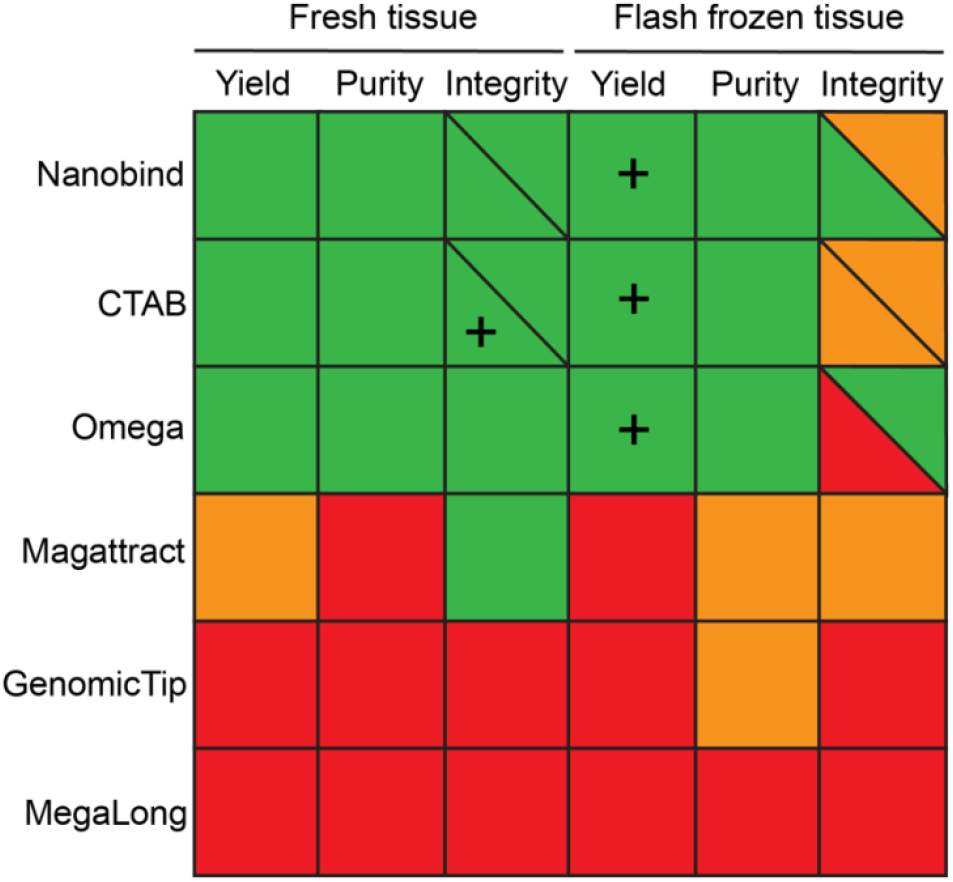
DNA extraction performance across methods. Thresholds for successful extraction were defined as: yield >1 µg total DNA (“+” indicates >10 µg across three replicates), purity with an A260/A280 ratio of 1.8–2.0, and integrity with the largest TapeStation peak >15 kb (“+” indicates largest TapeStation peak >60 kb across three replicates). Color coding: green = success in all replicates; orange = success in one to two replicates; red = no success. Extractions from five treatments (Nanobind fresh and frozen, CTAB fresh and frozen, Omega frozen) were submitted to the sequencing facility, where DNA integrity was confirmed on a Fragment Analyzer. Integrity values are shown as diagonals: lower left = TapeStation results from the extraction laboratory; upper right = Fragment Analyzer results from the sequencing center.

In contrast, the remaining three protocols—Nanobind, CTAB, and Omega—produced satisfactory DNA yield, purity, and integrity from fresh tissue, with CTAB yielding the highest integrity (Fig. 2; Table S1). As expected, the high-molecular-weight extraction methods (Nanobind and CTAB) generated DNA with greater integrity than the column-based Omega kit. Nonetheless, the Omega kit still produced relatively long fragments (approximately 18– 31 kb; Table S1), sufficient for several long-read sequencing applications. Interestingly, all three methods yielded substantially more DNA when using flash-frozen tissue, but this increase came at the expense of integrity, which was markedly reduced compared to extractions from fresh tissue (Fig. S1). Actually, the Omega kit did not meet our integrity threshold on flash-frozen material (largest TapeStation peak >15 kb). However, because the DNA profiles showed a clean and promising fragment distribution, we proceeded to submit these extracts to the sequencing center for further quality assessment. Some extracts, particularly CTAB from fresh tissue, showed relatively low A260/A230 ratios that may reflect residual carryover of contaminants (Table S1). However, because this metric was not consistently predictive of library preparation or sequencing success in our dataset, it was not used as a formal exclusion criterion.

To further assess potential improvements in DNA quality, we subjected extracts obtained with the two most promising column-free protocols (Nanobind and CTAB) from fresh tissue to additional clean-up steps using two commercial kits (Zymo and PowerClean; Fig. 3). Unexpectedly, the Zymo clean-up reduced the purity of Nanobind DNA, whereas CTAB-derived DNA remained stable after the same treatment. In both cases, the PowerClean protocol markedly degraded the DNA, rendering it unsuitable for long-read sequencing. However, DNA integrity after the Zymo clean-up remained sufficient for downstream long-read sequencing.

**Figure 3:**
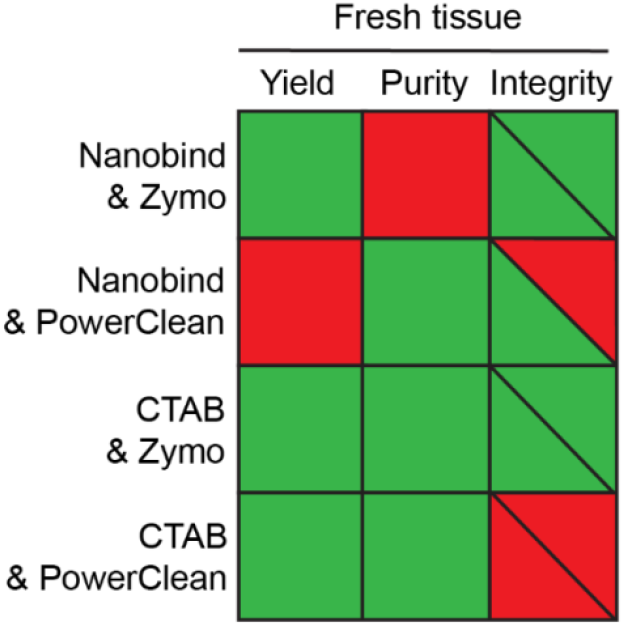
DNA quality metrics after additional clean-up step. Color coding: green = success. Red = no success. All four extractions were submitted to the sequencing facility, where DNA integrity was confirmed on a Fragment Analyzer. Integrity values are shown as diagonals: lower left = TapeStation results from the extraction laboratory; upper right = Fragment Analyzer results from the sequencing center.

### DNA extraction quality control, library preparation and PacBio sequencing at the sequencing center

Based on the quality control conducted at the DNA extraction laboratory in Dübendorf, 15 DNA extractions representing eight independent treatment conditions were selected for further quality control, library preparation, and sequencing at the GTF platform in Lausanne (Table 2). During the second round of quality control, substantial discrepancies in DNA integrity values were observed between instruments: the TapeStation (Eawag Dübendorf) and the Fragment Analyzer (GTF Lausanne) produced several inconsistent results (Fig. S2). These differences likely reflect both known platform-specific biases—since the TapeStation is known to slightly overestimate DNA fragment lengths—and most likely post-extraction DNA degradation (Fig. 2; Fig. 3).

For instance, Nanobind extractions from flash-frozen tissue displayed relatively high integrity when analyzed at the extraction laboratory but were found to be completely degraded upon re-evaluation at the sequencing facility (Fig. 2; Table 2; Fig. S2). Interestingly, this degradation was not observed for Nanobind extractions from fresh tissue, which retained their integrity across both analyses. A comparable pattern was found for CTAB extractions, where DNA from frozen tissue appeared substantially degraded at the sequencing center, while DNA from fresh tissue remained stable (Fig. S2). In contrast, no evidence of post-extraction degradation was detected for DNA obtained with the Omega kit; in fact, integrity values were relatively higher when assessed with the Fragment Analyzer (∼22-24 kb) compared to the TapeStation (∼13-14 kb), which was unexpected.

Following quality control at GTF, six samples (marked with asterisks in Table 2) were selected for library preparation. Because the initial run failed and used up the CTAB_fresh_Zymo1 library (see methods), only five libraries were ultimately sequenced. The second sequencing run was successful, yielding a total of 91 Gbp, which is in the range of expected output of a PacBio Revio cell (Table 3). One library (CTAB_frozen1) produced shorter reads, likely due to more extensive shearing compared to other samples, despite using the same shearing parameters. This resulted in reduced HiFi read length and overall yield. In contrast, the remaining libraries produced yields between 18 and 22.9 Gbp, with Omega_frozen1 achieving the highest output. A trade-off between HiFi read length and quality was observed, with shorter reads generally having higher quality. Nevertheless, even the longest reads (mean ∼13 kbp) maintained excellent quality (Q38). Overall, all five libraries were successfully sequenced, and read length distributions closely mirrored the respective library fragment sizes, indicating the absence of sequencing inhibitors.

**Table 3.** PacBio HiFi sequencing output on one Revio cell for each of the five libraries.

| Sample Name | HiFi Reads | HiFi Read Length (mean, bp) | HiFi Read Quality (median, QV) | HiFi Yield (Gbp) |
| --- | --- | --- | --- | --- |
| CTAB_fresh1 | 1,792,289 | 10,087 | Q41 | 18.08 |
| Nanobind_fresh1 | 1,493,698 | 12,371 | Q38 | 18.48 |
| Nanobind_fresh_Zymo1 | 1,469,733 | 13,051 | Q38 | 19.18 |
| Omega_frozen1 | 1,957,907 | 11,698 | Q39 | 22.91 |
| CTAB_frozen1 | 2,002,334 | 6,347 | Q47 | 12.71 |
| Not barcoded | 51,897 | 11,226 | Q30 | 0.58 |
| **Total** | **8,767,858** | **10,796** | **Q40** | **91.94** |

## Discussion

### Tissue preservation and extraction protocol shape extraction outcomes

Molluscs present well-known challenges for DNA extraction due to high levels of mucus and endogenous inhibitors (Adema, 2021). In this study, we evaluated different strategies for extracting high-molecular-weight (HMW) DNA from the freshwater mussel *Anodonta anatina*, a species for which we previously encountered difficulties in generating sufficient quality data for PacBio sequencing (Faust et al., 2025). While our results are based on a single species, we believe they provide valuable insights likely applicable to other unionids and possibly to other mollusc species.

We found that tissue preservation had a critical influence on the performance of the top three HMW DNA extraction protocols. This effect was evident both in terms of yield and integrity. First, yield differed markedly between preservation methods, with approximately ten times more DNA recovered from flash frozen tissue compared with fresh tissue. One possible explanation is that rapid freezing partially disrupted cellular membranes through intracellular ice-crystal formation (Muldrew and McGann, 1990). Therefore, flash freezing prior to extraction may have weakened tissue structure and improved lysis efficiency during DNA extraction, resulting in higher DNA recovery. While we did not explicitly test this mechanism, similar effects of increased DNA yield after flash freezing have been previously reported (Stark et al., 2020; Salis et al., 2025). For some samples, the NanoDrop DNA concentration measurements differed substantially from Qubit measurements, likely due to residual contaminants affecting absorbance-based quantification. We therefore considered NanoDrop values mainly as a complementary purity metric, while sequencing suitability was assessed primarily from Qubit yield, fragment integrity, and downstream library preparation success.

Furthermore, we observed clear differences in DNA integrity between fresh and flash-frozen tissue extracted using the Nanobind and CTAB protocols. While both protocols produced highquality and stable DNA from fresh tissue, DNA obtained from flash frozen tissue initially showed relatively high integrity at the DNA extraction laboratory in Dübendorf but exhibited strong post-extraction degradation when re-assessed at the sequencing center in Lausanne, in some cases declining from ∼18–28 kb or even >60 kb on the TapeStation to only ∼4 kb or less on the Fragment Analyzer. This occurred despite identical handling of all samples— storage at −20 °C after extraction and quality control, and transport on dry ice. The cause of this degradation remains unclear. One possibility is that residual compounds, such as DNases, remained active in extracts from frozen tissue, though this explanation remains speculative. Nevertheless, these findings demonstrate a clear advantage of extracting DNA from fresh tissue when feasible. We acknowledge, however, that this is often impractical—particularly in field-based or conservation genomic contexts—where optimized methods for preserved material remain indispensable (Reichel et al., 2026).

Among the flash frozen samples, the Omega kit consistently yielded DNA of moderate fragment length (∼13–14 kb on the TapeStation; ∼22-24 kb on the FragmentAnalyzer) and showed high reproducibility across replicates. Importantly, no post-extraction degradation was detected, suggesting that any residual compounds potentially responsible for degradation in the Nanobind and CTAB extractions were effectively removed by the Omega protocol. When applied to fresh tissue, the Omega kit also produced consistent results, yielding DNA of moderate fragment length (∼18–31 kb) and generally high purity. Together, these findings highlight the robustness and versatility of the Omega kit, which performed reliably across both fresh and flash frozen samples. An important limitation of this study is that all extractions were performed from foot tissue only. We chose foot tissue because it enables standardized sampling and, for fresh material, non-lethal biopsy while keeping the animal alive. However, tissue choice may itself influence extraction outcomes, because molluscan tissues are often rich in mucus, polysaccharides, and other compounds that can co-purify with DNA and inhibit downstream enzymatic reactions (Adema, 2021). Our results should therefore be interpreted as a comparison of extraction performance for *A. anatina* foot tissue, rather than as a general ranking of methods across all molluscan tissues.

### High-molecular weight DNA kits do not guarantee better results

In our study, three commercial HMW DNA kits (MagAttract, GenomicTip, Megalong) failed to yield usable DNA, from both fresh and flash frozen tissues. However, because performance may depend strongly on species, tissue type, and preservation method, these negative results should not be interpreted as a general failure of these kits across molluscs. In contrast, we successfully extracted high-quality DNA using a manual CTAB protocol, the Nanobind kit, and the Omega kit—though success depended strongly on tissue preservation. These results emphasize that dedicated HMW kits are not necessarily superior to manual or general-purpose protocols in molluscs and may require organism-specific optimization. Factors such as cost, processing time, and consistency should be considered when choosing a protocol.

Because mollusc DNA extractions are notoriously prone to inhibition by co-extracted contaminants, we tested additional post-extraction clean-up steps using PowerClean and Zymo kits to further remove residual inhibitors after standard DNA extraction. The PowerClean kit was unsuitable for genomic DNA, leading to severe degradation. In contrast, the Zymo kit preserved fragment length more effectively. However, clean-up did not provide clear added value, as libraries prepared from unpurified extractions (including from relatively dirty samples like CTAB_fresh1) sequenced successfully. In our case, additional clean-up of DNA extracts prior to PacBio library preparation did not provide additional benefits, although we cannot exclude that other clean-up strategies may be useful in different contexts.

We also note that the poor performance of some HMW extraction kits may partly reflect the specific properties of the tissue used here. Although foot tissue is practical for repeated and partly non-lethal sampling, kit performance observed here may not fully translate to other tissue types such as mantle, gill, or adductor muscle, which could differ in inhibitor load and tissue composition. Future work comparing multiple tissue types within the same individual would help disentangle protocol-specific from tissue-specific effects.

Ultimately, we generated high-quality PacBio data from five different DNA extraction and preservation combinations. Despite pooling all libraries on the same SMRT cell, we observed no evidence that inhibitors from one sample compromised sequencing success in others. Overall read yield was consistent across treatments, suggesting that inhibitor removal was generally successful, and that the selected protocols are robust enough for long-read sequencing applications in *A. anatina*.

### Recommendations for a new mollusc genome project

We showed that several HMW DNA extraction kits failed to produce DNA of sufficient quality for PacBio sequencing. Some protocols (PacBio Nanobind and CTAB) were successful, but their performance depended strongly on tissue preservation, leading to high-quality DNA only using fresh tissue. We would therefore recommend using one of these protocols if fresh tissue is available. It is also noteworthy that the manual CTAB protocol requires considerably longer hands-on time, while the Nanobind kit is comparatively rapid and user-friendly. Because we focused on *A. anatina*—a species notoriously difficult for DNA extraction (Faust et al., 2025)— rather than testing multiple species, we reduced confounding factors while generating results that provide a useful case study for mollusc genome projects, although their applicability across species, tissue types, and preservation methods should be interpreted cautiously.

In the present study, the Omega kit consistently produced DNA of sufficient quality for PacBio sequencing, especially for flash frozen tissue. The same kit has been successfully applied in the *Anodonta cygnea* genome project using flash frozen tissue (Faust et al., 2025). In another genome project conducted in our lab, we also obtained high-quality PacBio HiFi data from an RNAlater-preserved *A. exulcerata* sample stored at –80 °C using this kit – we however acknowledge that this is an anecdotal observation. Although ethanol-preserved tissues were not tested here, their widespread use makes them a relevant target for future evaluation with the Omega kit for PacBio HiFi sequencing.

We are aware that the Omega kit was not initally designed for long-read sequencing, and therefore fragment sizes are relatively modest. Hence, small adjustments to PacBio library preparation can - or should - be performed to increase the length of the resulting sequencing data. For example, in the present study, DNA was not sheared and was used directly in library preparation after the standard SRE step. In the above-mentioned *A. exulcerata* project, the sequencing platform further recommended a BluePippin purification step to remove small fragments (<10 kb) after PacBio library preparation. Nevertheless, such adjustments are straightforward, add little cost compared with months of DNA extraction optimization, and ensure effective sequencing output. In all cases, we recommend close contact and open communication with the experienced sequencing platform conducting library preparation and sequencing.

For new mollusc genome projects, it may often be most practical to begin with broadly reliable strategies rather than investing heavily at the outset in expensive HMW extraction kits and extensive protocol optimization. This is especially relevant because some of the dedicated HMW kits tested here (MagAttract, GenomicTip, and MegaLong) were among the most expensive options, yet did not yield DNA of sufficient quality, whereas the CTAB protocol and the Omega kit were substantially more cost-effective and performed better in the context of our study. Furthermore, the choice of DNA extraction method was strongly influenced by tissue preservation in *Anodonta anatina*. For fresh foot tissue, the PacBio Nanobind and manual CTAB protocols consistently produced the highest DNA integrity and are therefore the preferred options in this context. For flash frozen tissue, which is commonly used in field collections, the column-based Omega kit performed unexpectedly well, providing sufficient DNA quantity, purity, and fragment length for PacBio HiFi sequencing despite not being designed for HMW extraction. To our knowledge, this kit has also yielded high-quality PacBio data in three *Anodonta* species (Faust et al., 2025). The Omega kit does have limitations— adequate tissue input is needed to allow size selection, short fragments may require removal (e.g., via BluePippin), and it is not suitable when ultra-long reads are required. In such cases, Nanobind or CTAB remain better choices. Overall, our findings provide a practical decision framework for *A. anatina* and, more broadly, help guide initial protocol selection in other molluscan projects, although species-, tissue-, and preservation-specific validations should still be considered. Nanobind or CTAB are good options when high-quality fresh tissue is available, while the Omega kit represents an efficient and cost-effective alternative for flash frozen material.

## Acknowledgments

We thank the Canton of Zürich for providing the *Anodonta anatina* sampling permit, and Julie Conrads and Joana L. Santos for their assistance with field sampling. We are also grateful to Julien Marquis and Mélanie Dupasquier from the GTF platform at the University of Lausanne (UNIL) for their support with library preparation and PacBio sequencing. This work was supported by an Eawag Discretionary Fund Grant awarded to PGDF and AATW.

## Data availability

PacBio HiFi whole-genome sequencing data are available from the European Nucleotide Archive (ENA) under BioProject accession PRJEB105293. The dataset comprises five barcoded libraries and one set of unassigned reads generated from a single sequencing flow cell, all derived from the same BioSample (SAMEA120807678), with sequencing runs deposited as ERR16075080 and ERR16075215–ERR16075219.

## Author contributions

MG: Investigation, Supervision, Writing – review and editing. UL: Data curation, Investigation. PGDF: Funding acquisition, Writing – review and editing. AATW: Conceptualization, Funding acquisition, Project administration, Resources, Supervision, Visualization, Writing – original draft, Writing – review and editing.

## Notes

https://www.ebi.ac.uk/ena/browser/view/PRJEB105293

### Competing Interest Statement

The authors have declared no competing interest.

### Summary of Updates

This version includes a clean PDF with the PCI Genomics badge stating “Peer-reviewed and recommended by PCI Genomics”. The recommendation is available here: https://doi.org/10.24072/pci.genomics.100559

