## Supplementary figures and images for "Comparing DNA extraction methods for successful PacBio HiFi sequencing: a case study of the freshwater mussel *Anodonta anatina* (Bivalvia: Unionidae)"

### Figure S1

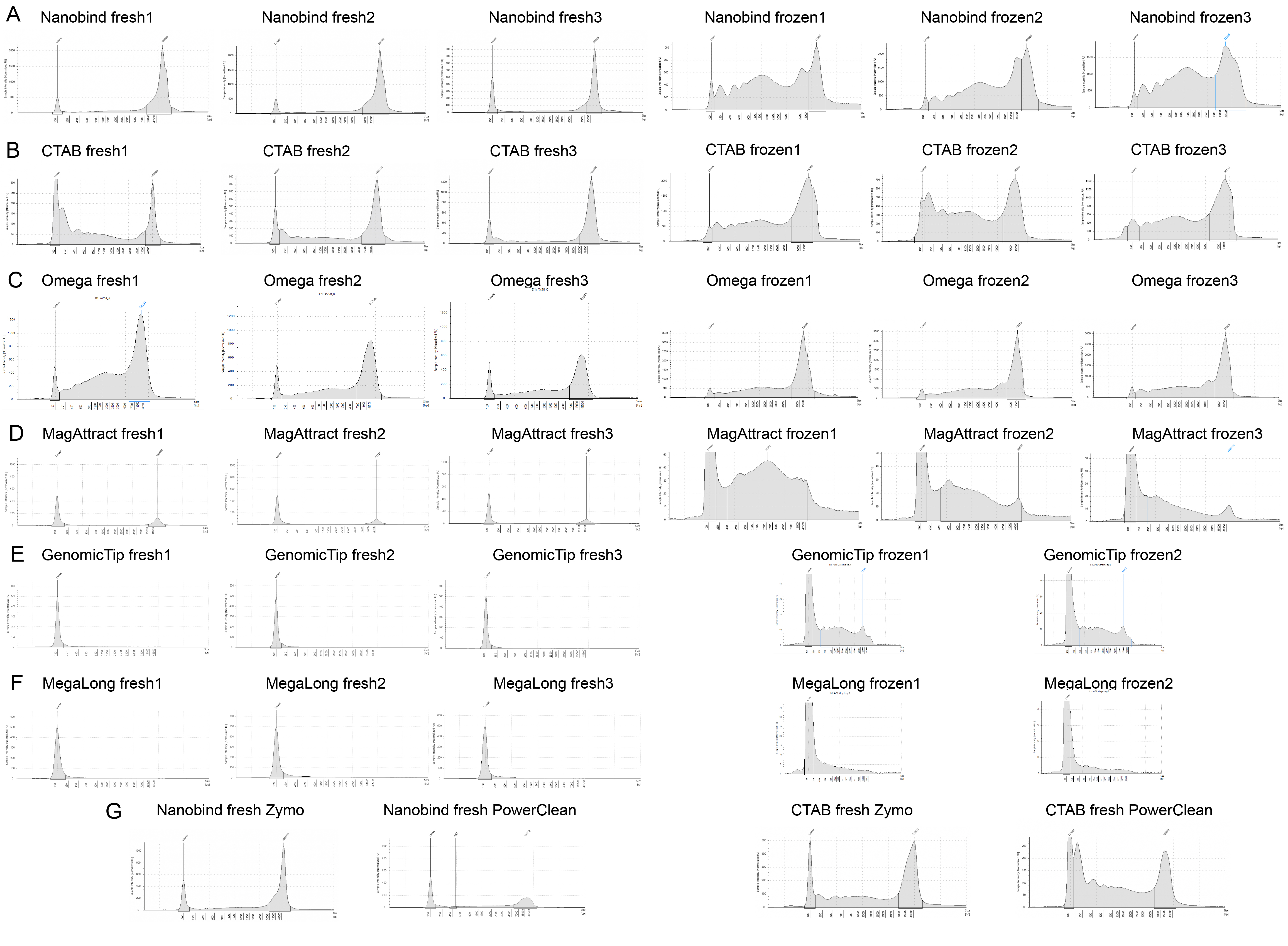

### Figure S2

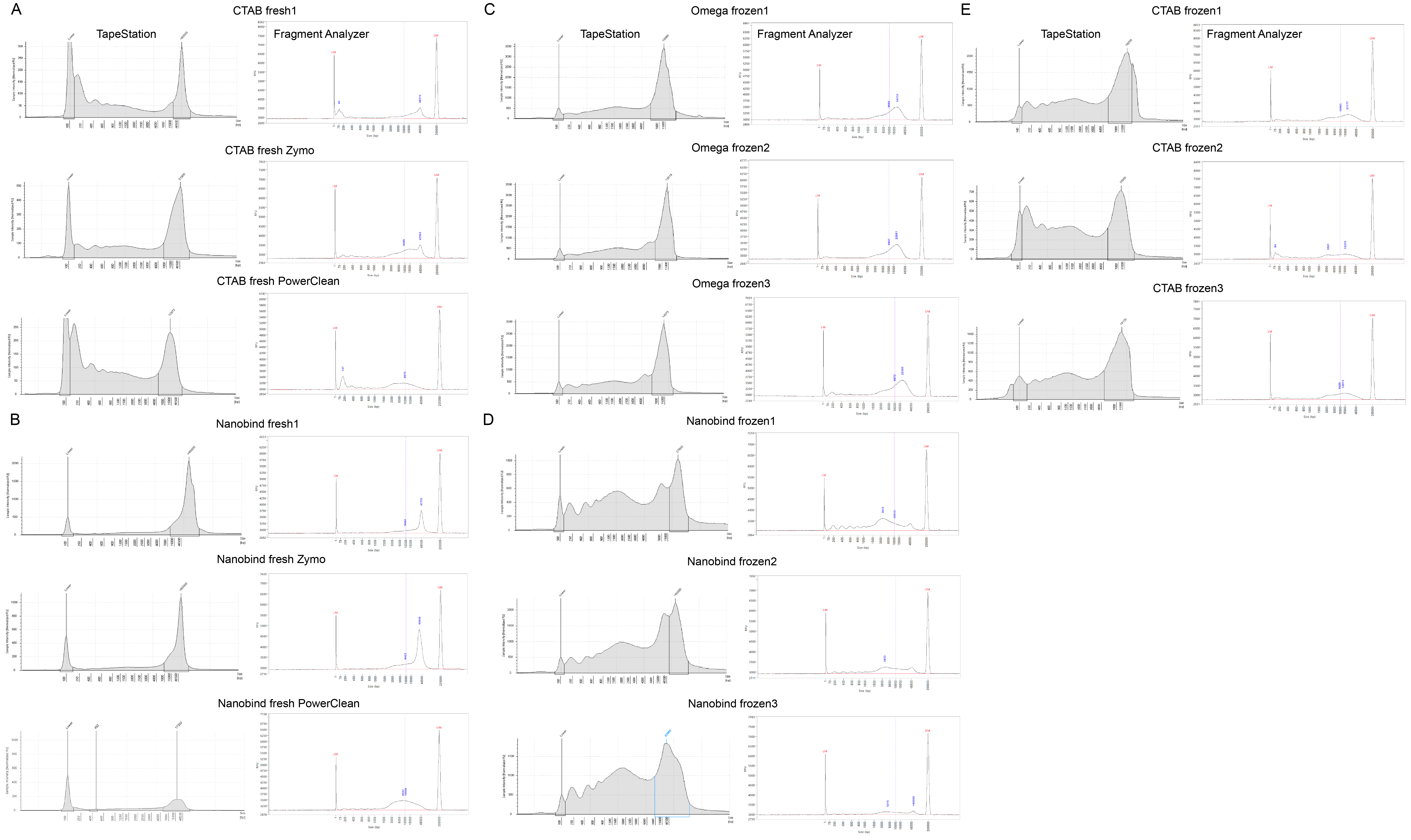
